# Circuit organization of the forelimb-related M2-to-M1 corticocortical pathway in the mouse

**DOI:** 10.64898/2026.02.03.703577

**Authors:** Louis Richevaux, Rita Fischer, Miraya Baid, Gordon M.G. Shepherd

## Abstract

Communication from secondary (M2, premotor) to primary (M1) motor cortex is implicated in forelimb motor control. We investigated the underlying synaptic circuits in this corticocortical pathway in male and female mice using cell-type-specific optogenetic-electrophysiology methods, focusing on identifying the cell-type-specific synaptic connections in the excitatory and feedforward inhibitory circuits impinging on cervically projecting M1 corticospinal neurons. In forelimb M1 brain slices, recordings from layer 5B corticospinal neurons during brief photostimulation of M2 axons showed strong monosynaptic excitatory currents that, although accompanied by potent feedforward inhibitory currents, were capable of evoking action potentials (APs) in most neurons. In contrast, responses in layer 2/3 pyramidal neurons were generally much weaker. Parvalbumin-expressing neurons (PV), particularly in deeper layers, showed direct excitation from M2 axons without feedforward inhibition, and could fire APs robustly. Somatostatin (SST) neurons received generally weak inputs, whereas VIP and Ndnf neurons received stronger excitation and inhibition from M2 axons. Corticospinal neurons received little or no local inhibition from Ndnf and VIP interneurons, but relatively strong soma-targeting PV and dendrite-targeting SST inhibitory inputs, as functionally imaged by laser-scanning synaptic input mapping (“sCRACM”). The domains of PV and SST inputs were partly overlapping around the corticospinal somata, but broader for PV and more vertical for SST inputs. Collectively, the results provide a working model for the cell-type-specific synaptic circuits of this “top-down” corticocortical pathway, organized around direct M2 excitation and PV-mediated inhibition of M1 corticospinal neurons.

**SIGNIFICANCE STATEMENT:** Cervically projecting corticospinal neurons in the primary motor cortex (M1) serve as the most direct conduits by which motor cortical activity reaches and influences spinal circuits controlling forelimb movements. Corticospinal activity is in turn influenced by inputs from multiple upstream areas. Here we studied the inputs from the secondary motor cortex (M2), a premotor-like area in the mouse, and characterized the patterns of synaptic connectivity formed by M2 axons onto multiple postsynaptic cell types in M1. The resulting “wiring diagram” suggests that these inter-areal circuits are configured to give premotor cortex privileged access to modulate M1 corticospinal output, through cell-type-specific connections and inhibitory mechanisms.

## INTRODUCTION

The forelimb-related subregion of primary motor cortex (M1) is a key node within a network of cortical areas that operate interactively to shape cortical output to subcortical circuits controlling movements of the hands and upper extremities. These areas are anatomically linked by inter-areal corticocortical axonal projections. The synaptic circuit connections within some of these pathways feeding into forelimb M1 is becoming better understood. For example, in the “bottom-up” direction, studies in rodents have characterized cell-type-specific circuits and in vivo dynamics in the corticocortical pathway linking the hand/forelimb-related subregions of primary somatosensory cortex (S1) and M1 (Yamawaki et al., 2021; Piña Novo et al., 2025).

The circuit organization of “top-down” corticocortical pathways linking forelimb-related premotor (secondary motor, M2) cortex to M1 is less well understood. Studies in rodents have identified various aspects, including anatomical projection patterns of axons, layer-related circuit features, and some of the excitatory connections of pyramidal neurons (Hira et al., 2013; Hooks et al., 2013; Ueta et al., 2013; Suter and Shepherd, 2015). However, many aspects remain incompletely characterized, including the connectivity of inhibitory interneurons and their roles in shaping M2-to-M1 communication. The extent to which patterns of corticocortical connectivity identified between other areas pertain to premotor-to-motor pathways is unclear. In vivo studies using multi-area recording methods are revealing M2-to-M1 inter-areal dynamics during diverse forms of behavior (Barthas and Kwan, 2017; Veuthey et al., 2020; Terada et al., 2022; Kristl et al., 2025; Saiki-Ishikawa et al., 2025). Interpreting these dynamics in terms of underlying cellular mechanisms will entail elucidation of M2-to-M1 synaptic connectivity.

Here, to characterize the synaptic circuit organization of the forelimb-related M2-to-M1 corticocortical pathway in greater detail, we used an approach combining optogenetic-electrophysiological methods together with multiple interneuron-specific mouse lines and retrograde labeling of corticospinal neurons (Yamawaki et al., 2016). We evaluated the cell-type-specific synaptic connectivity between identified pre- and postsynaptic neurons, with a focus on direct excitation and feedforward inhibition to M1 corticospinal neurons. The results collectively provide a detailed working framework for this corticocortical pathway, which appears specifically configured to support direct excitation of M1 corticospinal neurons by M2 axons, accompanied by feedforward inhibition mediated by PV interneurons.

## METHODS

### Animals

The animal studies were approved by the Northwestern University Animal Care and Use Committee. Experiments were performed on wild-type (WT) C57BL/6 mice (Jax #000664) and transgenic lines, including: PV-Cre (Jax #017320) (Hippenmeyer et al., 2005), SST-Cre (Jax #013044) (Taniguchi et al., 2011), VIP-Cre (Jax #0311628) (Taniguchi et al., 2011), and Ndnf-Cre (Jax #028536) (Tasic et al., 2016). These Cre driver lines were crossed with reporter lines Ai14 (Jax #007908) (Madisen et al., 2010), Ai32 (Jax #012569) (Madisen et al., 2012), or Ai203 (Jax #037939) (Bounds et al., 2023) to express red fluorescent protein tdTomato, channelrhodopsin-2 (ChR2) fused with GFP, or soma-targeted ChroME with mRuby, respectively, in the different types of interneurons. All lines were maintained on a C57BL/6 background. Male and female animals were used in approximately equal numbers. Mice were housed with a 12 hr light/dark cycle. Mice were 5-6 weeks of age at the time of surgery and were used for experiments 3-4 weeks later.

### Viruses

Adeno-associated viruses (AAV) used in this study included AAV1.CamKII.ChR2(H134R)-eYFP (Addgene #26969-AAV1) (Lee et al., 2010), used to anterogradely express ChR2 in M2 afferents to M1, and AAVretro.CAG.tdTomato (Addgene #59462-AAVrg) or AAVretro.CAG.GFP (Addgene #37825-AAVrg) (Tervo et al., 2016), used to retrogradely express tdTomato or GFP in corticospinal neurons after injection in the spinal cord. Viruses were stored at -80°C.

### Injections

Viral vectors were stereotaxically injected as previously described (Yamawaki et al., 2021). Mice were anesthetized with isoflurane and positioned in a stereotaxic frame with a thermal support. Prior to surgery, animals were administered buprenorphine (1 mg/kg) and bupivacaine (2 mg/kg) subcutaneously for analgesic coverage. Craniotomies were opened above the intended sites of cortical injections in the right hemisphere. For spinal cord injections, a laminectomy was performed at cervical level 6 (C6) to access the left side of the cord for injections. Injection pipettes were pulled from borosilicate glass capillaries and beveled.

Premotor cortex (M2) injection coordinates were (relative to bregma) 2.0 mm antero-posterior and 1.2 mm lateral, based on the presence of retrogradely labeled corticospinal neurons projecting to C6 (Hausmann et al., 2022). In the cortex, injections (50 nL each) were made at depths of -0.3, -0.6, and -0.9 mm. In the spinal cord, a single injection (300 nL) was made at the C6 level.

After injections were complete, the surgical wound was closed using non-absorbable sutures, and mice were injected with meloxicam (20 mg/kg) and returned to a clean cage. Meloxicam was again administered 24 h after.

### Slice electrophysiology

Mice were euthanized 3-4 weeks after injection and 300 µm-thick coronal brain slices were prepared using a vibratome in ice-cold choline-based solution (containing in mM: 25 NaHCO_3_, 1.25 NaH_2_PO_4_, 2.5 KCl, 0.5 CaCl_2_, 7 MgCl_2_, 110 choline chloride, 11.6 sodium L-ascorbate, and 3.1 sodium pyruvate). Slices were transferred to artificial cerebrospinal fluid (ACSF; composition, in mM: 127 NaCl, 25 D-glucose, 2.5 KCl, 1 MgCl_2_, 2 CaCl_2_, and 1.25 NaH_2_PO_4_) for 30 min at 34°C and then 1 h or more at room temperature (∼22 °C) before recording.

Whole-cell recordings were performed using an upright microscope equipped with gradient contrast and epifluorescence optics. Pipettes (∼2.5–4 MΩ) were filled with cesium-based internal solution containing (in mM): 128 cesium methanesulfonate, 10 HEPES, 10 phosphocreatine, 4 MgCl_2_, 4 ATP, 0.4 GTP, 3 ascorbate, 1 QX314, and 1 EGTA. To record action potentials, we used potassium-based internal solution containing (in mM): 128 potassium methanesulfonate, 10 HEPES, 10 phosphocreatine, 4 MgCl_2_, 4 ATP, 0.4 GTP, 3 ascorbate, 1 EGTA. Internal solutions were balanced at pH 7.25, 290–295 mOsm. ACSF, oxygenated with 95% O_2_/5% CO_2_, was perfused in a closed loop, and the recording temperature was maintained at 34 °C. To record monosynaptic responses, we added TTX (1 µM, Tocris Bioscience) and 4-AP (100 µM, Sigma-Aldrich) to the bath solution (Petreanu et al., 2009); to record polysynaptic responses (feedforward inhibition), these drugs were omitted. We also routinely blocked NMDA-receptor currents (CPP, 5 µM, Tocris Bioscience) to avoid non-linear dendritic responses. In control experiments, excitatory inputs were abolished after application of an AMPA receptor antagonist (NBQX, 10 μM; n = 2 neurons), confirming that these currents were carried mainly by fast ionotropic glutamatergic synaptic transmission (**Figure 2B**). Signals were amplified with an Axon Multiclamp 700B (Molecular Devices), filtered at 4 kHz, and sampled at 40 kHz. Data acquisition was controlled by Ephus software (Suter et al., 2010).

### Wide-field photostimulation

Photostimuli were delivered via the epifluorescence pathway as 5 ms square-pulses of blue LED light (473 nm wavelength, M470L2 source, LEDD1B driver, Thorlabs, Inc., excitation filter HQ480/20, 4x magnification). Stimuli were repeated 5 times with 30 s intervals. Responses were recorded in voltage clamp mode, with the command potential set to -70 mV for recording excitatory postsynaptic currents or potentials (EPSC/Ps), and +10 mV for inhibitory postsynaptic currents (IPSCs).

### Subcellular ChR2-assisted circuit mapping (sCRACM) and laser-scanning photostimulation (LSPS)

Inputs were mapped by sCRACM (Petreanu et al., 2009) using a laser-scanning system equipped with a 473 nm laser (MLL-FN473-50mW, Opto Engine LLC). Under 4x magnification, the location of the soma was marked, and a 30-by-10 stimulation grid was overlaid onto the bright-field image. Stimulation sites were equally spaced every 50 µm. The grid was horizontally centered on the soma and vertically aligned to the pia. The height and width of the grid (1500-by-500 µm) covered all layers and spanned the dendritic arbors of corticospinal neurons, as previously described (Suter and Shepherd, 2015). For each neuron recorded, laser power was adjusted to elicit ∼300 pA peak responses for stimulation near the soma. Stimulation sites were visited in a pseudorandom sequence, with an inter-site interval of 1 s. Each neuron was mapped 2–3 times.

For LSPS mapping of inputs in Ai203-crossed mice, we used the same approach described above for sCRACM, but with 16-by-16 grids with 100 µm spacing aligned on the pia. In the Ai203 line, all neurons with Cre express nuclear-targeted mRuby, and a subset also expresses the st-ChroME construct (Bounds et al., 2023). In control experiments, we recorded excitation profiles of channelrhodopsin expressing neurons, in the presence of NBQX and gabazine, to determine suitable laser power for reliably evoking action potentials and confirm soma-targeted expression.

### Slice histology and imaging

Brain sections were cut, mounted, and imaged by confocal microscopy as previously described (Yamawaki et al., 2021; Hausmann et al., 2022). Laminar profiles of fluorescence intensity were measured using ImageJ software. Forelimb M1 and M2 areas were localized based on the presence of corticospinal neurons, as well as anatomical landmarks and reference to brain atlases (Allen Brain Atlas - http://mouse.brain-map.org). Cortical layers in forelimb M1 were identified based on well-established cytoarchitectonic features and layer-defining labeling patterns determined in previous studies (Yamawaki et al., 2021). Cortical layers in forelimb M2 were identified based on similar features and labeling patterns. In addition to using corticospinal labeling to identify layer 5B, we used layer-specific gene expression patterns and reference atlases. Unlike more lateral cortical areas on the dorsal surface of the mouse brain, and as previously described for vibrissal M1 (Hooks et al., 2011), the cortical layers of forelimb M2 display a geometric transformation commonly associated with cortical gyri and flexures, wherein superficial layers are thinner and wider and deeper layers are narrower and taller. Thus, layer 5A is relatively more superficial in M2 compared to M1.

### Data analysis

Electrophysiological recordings were analyzed off-line using MATLAB. All postsynaptic responses were averaged over multiple trials and baseline subtracted (50 ms prestimulus). For recordings in voltage-clamp mode, input strength was measured by averaging the amplitude of the postsynaptic response over 100 ms poststimulus for recordings. For current-clamp recordings, action potentials were counted over 50 ms poststimulus. Firing rate as a function of current step amplitude was calculated over a duration of 1 s in steps of 50 pA increments up to 500 pA. Example traces of recordings in voltage-clamp mode show single neuron average traces (5 trials), and those of action potential individual traces. Group-averaged traces show mean ± SEM of every neuron’s average trace.

Cross-correlation analysis of LSPS mapping results was performed as follows. First, for each neuron, we averaged along the rows of the 2-dimensional input map to obtain a 1-dimensional vector representing the vertical profile of input. We then calculated the cross-correlations between all neurons’ vertical profiles. For each pairwise cross-correlation, we found the peak value and the corresponding lag (i.e., the distance) at the peak. In other words, in this step we determined how much the vertical profiles needed to be shifted vertically (up toward the pia or down toward the white matter) for optimal alignment. We then constructed a matrix with these lag values, sorted the rows and columns by the soma depth in the cortex, and plotted these as a pseudocolored image. Thus, the lags (shifts) would be expected to be non-zero and directly proportional to soma depth if input map structure is solely determined by the targeting of presynaptic axon terminals to particular postsynaptic somatodendritic domains (e.g. soma- or dendrite-targeting). In contrast, lags would be expected to be zero if input maps are determined solely by the laminar location of the presynaptic axons.

Anatomical analysis of labeling intensity in fluorescence images was performed using the Plot Profile function in Image J. Laminar profiles of labeling intensity were obtained by measuring across the cortical depth (i.e., spanning the pia to the white matter underlying L6). Data were subsequently normalized, resampled, and averaged using standard MATLAB functions.

### Statistical Analysis

Central tendency is expressed as median ± median absolute deviation (MAD) unless stated otherwise. Inhibition-excitation (IE) ratio central tendency is expressed as geometric mean, and dispersion as the following range: (from geometric mean ÷ geometric standard deviation (SD) to geometric mean × geometric SD). Mann-Whitney U test was used to compare two unpaired cell groups. The Kruskal-Wallis test followed by multiple comparisons correction using the Dunn-Sidak test was used to compare more than two groups. Where specified, one-way ANOVA was used to compare repeated measures in two groups. In all cases, statistical significance was set at *p* = 0.05.

## RESULTS

### M2 projections to M1 ramify in corticospinal-associated layers

As an initial step toward characterizing the synaptic connectivity from M2 to M1 neurons including corticospinal neurons, we imaged the corticocortical axonal projection in this pathway. First, as the presence of cervical-projecting corticospinal neurons is a landmark of both forelimb M1 and forelimb M2 (Neafsey and Sievert, 1982; Nudo and Masterton, 1990), we injected the spinal cord at cervical level C6 with a retrograde virus carrying a fluorescent protein (AAVrg.CAG.GFP or tdTomato; henceforth referred to as AAVretro-FP), thereby precisely identifying and localizing these two areas (Hausmann et al., 2022) (**Figure 1A**). Of note, the more posterior and larger of these two patches of corticospinal labeling spans not only forelimb M1 but extends to hand/forelimb S1, with forelimb M1 situated anteromedially within this patch, as described previously in detail (Ueno et al., 2018; Yamawaki et al., 2021). We then used retrograde labeling from M1 to localize the M2 origin of this pathway. In wild-type mice, we stereotaxically injected M1 with another AAVretro-FP carrying a different fluorescent protein to label the M1-projecting neurons (**Figure 1B**). In forelimb M2, these neurons were broadly distributed across layers but mainly located across the middle layers, particularly in layer 5.

**Figure 1.**
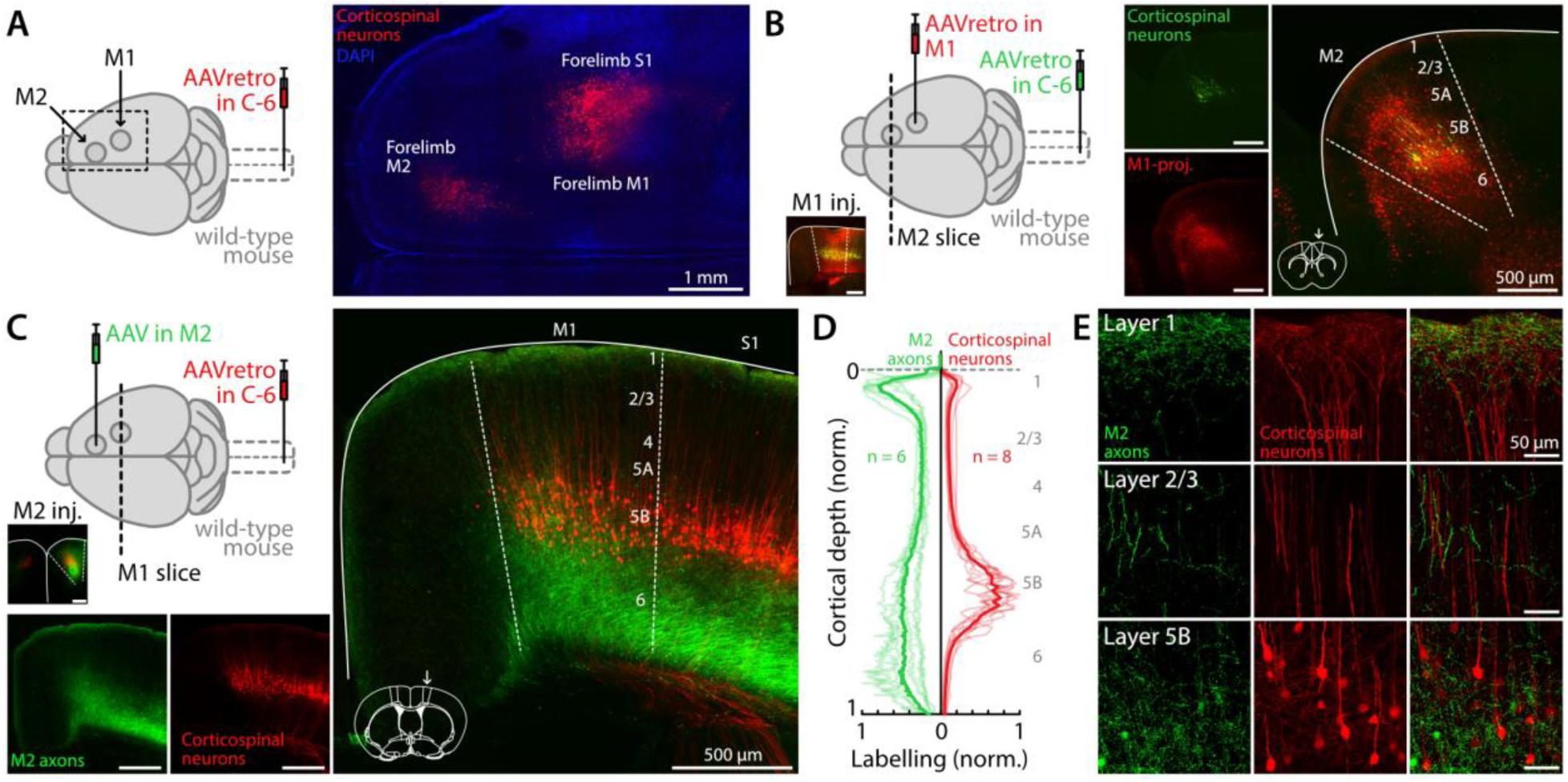
M2 projections to M1 ramify in corticospinal-associated layers. **A**, Left: Schematic of the labeling strategy. Retrograde AAV was injected in the spinal cord (cervical level C6) to label corticospinal neurons. Right: Fluorescence image of a superficial horizontal brain slice, showing labeling of corticospinal neurons in forelimb M2, M1, and S1 (red). **B,** Left: Schematic of the labeling strategy. Retrograde AAV was injected in the spinal cord (cervical level C6) to label corticospinal neurons, and in M1 to label M1-projecting neurons in M2. M1 injection site is shown in inset. Right: Fluorescence image of an M2 coronal brain slice, showing labeling of M1-projecting corticocortical neurons (red) and corticospinal neurons (green). Coronal slice schematic adapted from Paxinos and Franklin atlas (2013). **C,** Left: Anterograde AAV was injected in M2, and retrograde AAV was injected in the spinal cord. M2 injection site is shown in inset. Bottom and right: Fluorescence image showing labeled M2 (green) axons and corticospinal neurons (red) in an M1 coronal slice. **D,** Fluorescence profile of M2 afferents (green, n = 6 slices, 6 animals) and corticospinal neurons (red, n = 8 slices, 7 animals) in M1, normalized vertically to the pia and whiter matter border, and to the minimum/maximum fluorescence intensity. **E,** Images show 60x views of labeling in M1 layers 1, 2/3, and 5B, separately for M2 axons (left column), corticospinal neurons/dendrites (middle column), and both (right column).

To image the axonal projection from M2 to M1, we injected M2 with an anterograde virus carrying a fluorescent protein (AAV1.CAG.GFP; henceforth AAV-FP). Again, we also injected the spinal cord with AAVretro-FP (**Figure 1C**), for positive identification and localization of forelimb M2 and M1. Targeting of the M2 injection site was confirmed by the presence of green fluorescent labeling surrounding red-labeled corticospinal neurons. In M1, the labeled M2 afferents were distributed as a broad band in the deeper layers (layers 5B/6) and a narrow band in superficial layer 1, separated by an upper-layer zone of lower-density labeling spanning layer 2/3 (**Figure 1C**). The labeled corticospinal neurons were located in layer 5B. The laminar profiles of M2 axon labeling overlapped with those of the labeled corticospinal neurons and their dendrites, with higher labeling intensity in layers 1 and 5B/6 (**Figure 1D**). Higher-magnification views showed the axonal and dendritic labeling patterns in greater detail (**Figure 1E**). Thus, M2 afferents appear to overlap anatomically with M1 corticospinal neurons, the dendrites of which have high density in basal/perisomatic and apical tuft regions (Suter and Shepherd, 2015).

### M2 input strongly excites M1 corticospinal neurons, with strong feedforward inhibition

To characterize the excitatory-to-excitatory (E→E) connections in the M2→M1 pathway, we injected M2 with an anterograde virus carrying ChR2 (AAV1.CamKII.ChR2(H134R)-eYFP; henceforth AAV-ChR2) (**Figure 2A**). We measured the strength of monosynaptic excitatory currents evoked by wide-field photostimulation of these axons with blue light (473 nm, 10 mW, 5 ms) in forelimb M1 retrogradely labeled corticospinal neurons and (unlabeled) layer 2/3 (L2/3) pyramidal neurons (**Figure 2A, B**). Pharmacological conditions were set (by addition of TTX and 4AP to the bath solution) to restrict responses to direct, monosynaptic input (Petreanu et al., 2009)(**Methods**). Optogenetic activation of M2 afferents evoked excitatory postsynaptic currents (EPSCs) in corticospinal and L2/3 pyramidal neurons. On average, responses were several-fold stronger in corticospinal compared to L2/3 pyramidal neurons (103.6 ± 44.2 pA vs 22.5 ± 15.4 pA, median ± MAD; *p* = 6×10^-4^, Mann-Whitney U test; 18 corticospinal and 17 L2/3 pyramidal neurons, from 4 mice) (**Figure 2C**). These results thus indicate that M2 axons provide relatively strong monosynaptic excitation (E→E) to M1 corticospinal neurons (**Figure 2D**).

**Figure 2.**
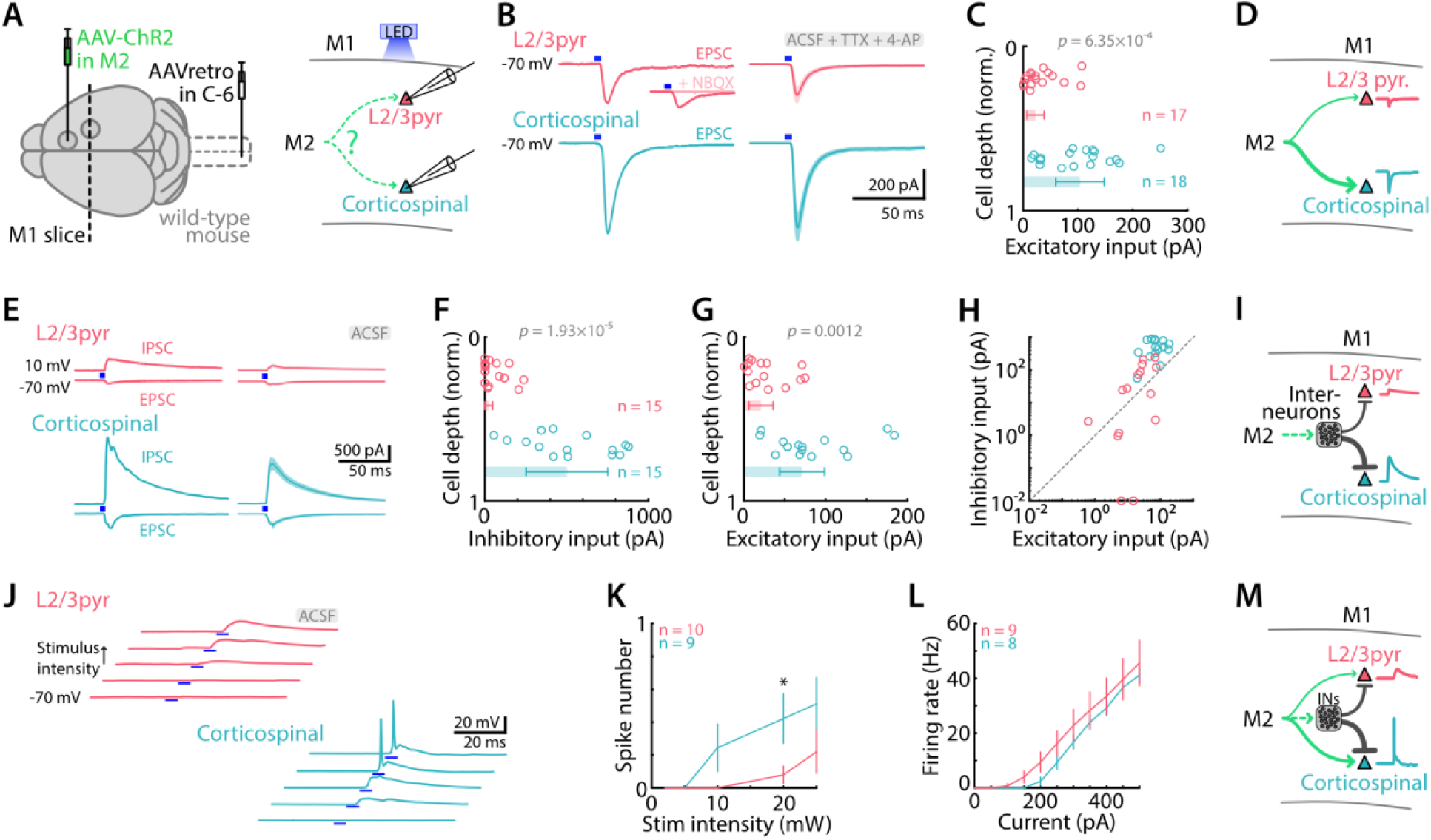
M2→M1 corticocortical input strongly excites corticospinal neurons, with feedforward inhibition. **A**, Schematic depiction of the labeling and recording paradigms. Left: Anterograde AAV was injected in M2 to induce ChR2 expression in M1-projecting neurons, and corticospinal neurons were labeled by retrograde AAV injected in the spinal cord. Right: M2 afferents were stimulated with blue light LED while M1 L2/3 pyramidal and corticospinal neurons responses were recorded. **B,** Example (left) and average (right, mean ± SEM) monosynaptic excitatory responses of L2/3 (top, red), and corticospinal neurons (bottom, blue) to M2 stimulation (5 ms, 10 mW). Recordings were performed in artificial cerebrospinal fluid (ACSF) with TTX (1 µM) and 4-AP (100 µM). Blue rectangles indicate LED stimulation. Inset shows absence of response in presence of AMPA receptor antagonist NBQX (10 µM). **C,** Excitatory input to L2/3 and corticospinal neurons in M1, calculated as the average response amplitude over 100 ms post-stimulus and plotted as a function of normalized cortical depth of the soma (0 = pia, 1 = border of L6 with underlying white matter). Bars show the median ± MAD. **D,** Schematic depiction of the excitatory connectivity pattern. **E,** Example (left) and average (right, mean ± SEM) traces showing excitatory (−70 mV) and inhibitory (10 mV) responses to photostimulation of M2 axons, recorded in individual corticospinal (bottom, blue) and L2/3 pyramidal neurons (top, red), in permissive (control ACSF) conditions. **F,** Amplitudes of inhibitory responses plotted against normalized cortical depth of the soma. **G,** Same for excitatory responses **H,** Amplitudes of excitatory and inhibitory responses recorded for each neuron. **I,** Schematic depiction of the feedforward inhibition connectivity pattern. **J,** Example traces of excitatory postsynaptic potentials evoked by M2 axons photostimulation with increasing light intensity (2, 5, 10, 20, 25 mW) in individual L2/3 (top, red) and corticospinal neurons (bottom, blue). Higher intensity generated action potentials in corticospinal neurons. **K,** Average number of evoked action potentials (spike number) as a function of stimulus intensity (mean ± SEM) for L2/3 and corticospinal neurons. Asterisks denote *p* value below 0.05 (one-way ANOVA). **L,** Frequency-current plot, showing the average (mean ± SEM) firing rate (number of action potentials per 1-s stimulus) as a function of current step amplitude for L2/3 (red) and corticospinal neurons (blue). **M,** Schematic depiction of the connectivity pattern.

To measure feedforward inhibitory (E→I→E) currents in the same two classes of postsynaptic pyramidal neurons (**Methods**), we repeated the same experiments but with pharmacological conditions set to allow rather than prevent polysynaptic responses (i.e., by omitting TTX/4AP). We used Cs^+^-based intracellular solution, setting the command potential at - 70 mV or 10 mV to record EPSCs or inhibitory postsynaptic currents (IPSCs) in voltage clamp mode. Photostimulation (10 mW, 5 ms) of M2 axons elicited both EPSCs and IPSCs in corticospinal and L2/3 pyramidal neurons (**Figure 2E**). The evoked IPSCs were nearly 20-fold stronger in corticospinal than in L2/3 pyramidal neurons (503.2 ± 259.0 pA vs 24.4 ± 24.4 pA, *p* = 1.9×10^-5^, 15 corticospinal and 15 L2/3 pyramidal neurons, from 7 mice) (**Figure 2F**). In the same neurons, the evoked EPSCs were again several-fold stronger in corticospinal compared to L2/3 pyramidal neurons (71.4 ± 27.5 pA vs 21.0 ± 14.7 pA, *p* = 0.0012) (**Figure 2G, H**). Inhibition-excitation (IE) ratios were accordingly much higher for corticospinal compared to L2/3 pyramidal neurons (geometric mean and SD: 6.09 (2.75 to 13.48) vs 0.53 (0.03 to 10.04), *p* = 0.0037). These results thus indicate that the strong direct excitation from M2 axons to M1 corticospinal neurons is accompanied by strong feedforward (polysynaptic, E→I→E) inhibition (**Figure 2I**). Given that M2-driven direct excitation and feedforward inhibition to corticospinal neurons are both strong, but with relatively more inhibition, we wanted to assess the net effect of activating M2 inputs on postsynaptic excitability, including the ability to drive action potentials. To do so, we recorded the responses of these neurons in current clamp mode with a K^+^-based internal solution, allowing the generation of action potentials. Stimulation of M2 afferents with lower light intensities evoked subthreshold excitatory postsynaptic potentials (EPSPs), while stimulation with higher light intensities evoked action potentials (**Figure 2J**). However, as shown by plotting spike number as a function of stimulus intensity, M2 inputs evoked spikes at lower intensities in corticospinal neurons than in L2/3 pyramidal neurons (*p* = 0.014 at 20mW, one-way ANOVA, 10 L2/3 and 9 corticospinal neurons, from 4 animals) (**Figure 2K**). The greater suprathreshold excitability of corticospinal neurons to M2 input was not ascribable to differences in the frequency-current relationships of the two cell types, as these were similar (*p* = 0.66, one-way ANOVA, 9 L2/3 and 8 corticospinal neurons, from 4 animals) (**Figure 2L**). These results thus indicate that the net effect of M2 input is strongly excitatory in M1 corticospinal neurons (**Figure 2M**).

### M2 input also excites L2/3 pyramidal neurons, across their dendritic domains

The strong M2 excitation of M1 corticospinal neurons accords with previous dendritic-level mapping studies showing M2 inputs to both perisomatic dendrites in deeper layers and to apical tuft dendrites in layer 1 (Suter and Shepherd, 2015). Although the current study focuses primarily on inputs to M1 corticospinal neurons, the M2 excitation of M1 L2/3 pyramidal neurons is also of interest, both for the question of the extent to which the input reported above is onto apical dendrites in layer 1 versus perisomatic dendrites in layer 2/3, and because L2/3 pyramidal neurons are the primary source of excitatory input to corticospinal neurons (Weiler et al., 2008; Anderson et al., 2010). We therefore used sCRACM, a mapping method based on laser-scanning photostimulation (LSPS) across a grid of focal stimulation sites, to map the locations of evoked input from M2 axon presynaptic terminals onto L2/3 pyramidal neuron dendrites (Methods) (**Figure 3A**). These results showed that while neurons located in superficial layer 2/3 received M2 input at layer 1 dendrites in addition to their perisomatic dendrites, there was a significant drop-off of layer 1 inputs for deeper neurons (**Figure 3B-E**). Indeed, layer 1 inputs decreased while inputs from other layers increased (**Figure 3E**, top). However, this loss of apical inputs for deeper cells did not change the overall input as LED stimulation evoked responses of similar amplitude in upper L2/3 pyramidal neurons compared to deeper neurons (**Figure 3E**, bottom). This pattern, similar to that found for corticospinal neurons previously (Suter and Shepherd, 2015), could be due to distance-dependent electrotonic attenuation (Williams and Mitchell, 2008), or to selectivity in the synaptic connectivity due to other factors, or a combination thereof.

**Figure 3.**
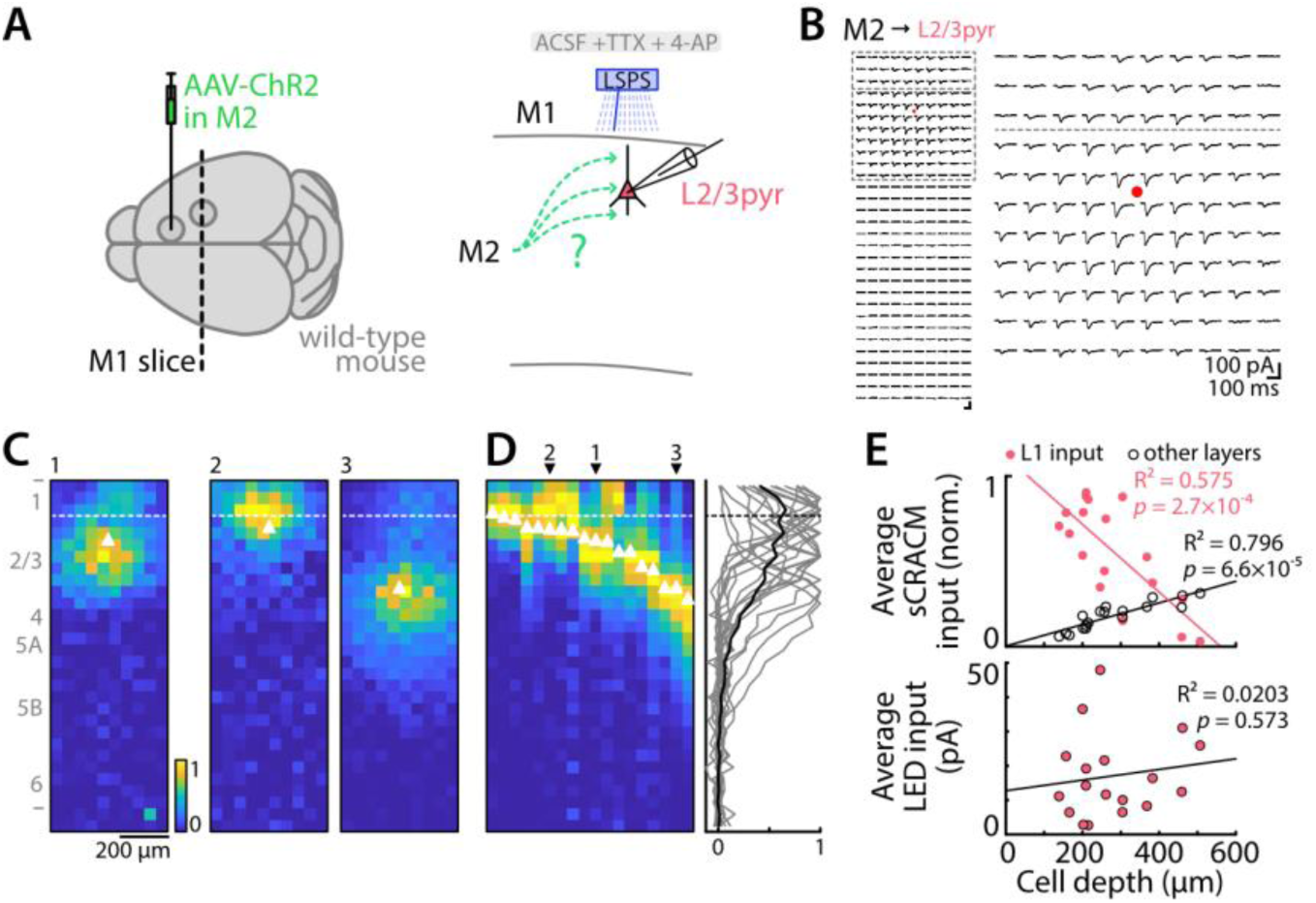
sCRACM-mapping of excitatory inputs from M2 axons to M1 L2/3 pyramidal neuron dendrites. **A**, Schematic depiction of the labeling and recording paradigms. Left: AAV-ChR2 was injected in M2. Right: Laser scanning photostimulation (LSPS) was used to map the dendritic locations of M2 presynaptic terminals across the dendrites of the recorded M1 L2/3 pyramidal neuron under sCRACM recording conditions (Methods). **B,** Left: Example traces from an LSPS input map, showing responses evoked by focal laser stimulation in a 10×30 grid pattern (see Methods). Soma is indicated by a red point. Right: Enlarged view of perisomatic and apical regions. Horizontal dashed line: border between L1 and L2/3. **C,** LSPS maps of peak-normalized input strength for three example neurons (neuron 1 is from panel B traces). Soma is indicated by white triangle. Horizontal dashed line: border between L1 and L2/3. **D,** Horizontal average projection of each neuron’s input map sorted by soma depth (left). Numbers and arrowheads on the top indicate profiles of neurons shown in C. Right: Corresponding individual (thin lines) and overall average (thick line) vertical profiles. **E,** Top: Average normalized responses evoked by LSPS photostimulation at sites in L1 (pink dots) and at sites across all other layers (black circles), as a function of soma depth. Bottom: Average responses evoked by wide-field LED stimulation (5 ms, 25 mW), as a function of soma depth. Lines: linear regressions.

### M2 input strongly excites M1 PV interneurons without feed-forward inhibition

The strong feedforward inhibitory responses generated in M1 excitatory neurons (E→I→E) indicate strong engagement of M1 GABAergic interneurons by activation of M2 axons (E→I). Therefore, we next examined the relative strength of M2 excitatory input to different types of M1 interneurons. To do so, we performed the same experiments described above, but with recordings targeted to fluorescently labeled neurons in brain slices prepared from mice expressing tdTomato in different interneuron lines. We first focused on PV interneurons (PV-Cre x Ai14, **Figure 4A, B**), as these constitute a major class of cortical interneurons and are potential targets of corticocortical input (Muñoz-Castañeda et al., 2021; Naskar et al., 2021; Atsumi et al., 2023). PV interneurons were recorded across the full range of their distribution throughout the depth of the cortex (i.e., across all layers where found) (**Figure 4C, D**). L2/3 pyramidal neurons were also recorded for comparison (**Figure 4C**), rather than corticospinal neurons (which would have entailed an additional color channel and injection). For each neuron, we first recorded excitatory responses to M2 axon stimulation in monosynaptic restrictive conditions (i.e., addition of TTX/4AP to the bath solution), and analyzed these as described above for the recorded excitatory neurons. On average, L2/3 pyramidal neurons and PV interneurons responded with EPSC of similar strength to M2 input photostimulation (26.3 ± 17.0 pA vs 22.7 ± 9.8 pA, *p =* 0.72, 12 L2/3 pyramidal neurons, 23 PV interneurons, from 4 animals). However, EPSC amplitudes were greater for PV interneurons located in deeper layers than upper layers (32.9 ± 11.1 pA vs 16.4 ± 7.2 pA, *p* = 0.006, 14 neurons in the upper half of the cortex, 9 neurons in the lower half) (**Figure 4E-G**). These results thus demonstrate moderately strong monosynaptic input from M2 axons to PV interneurons in M1, particularly in deeper layers.

**Figure 4.**
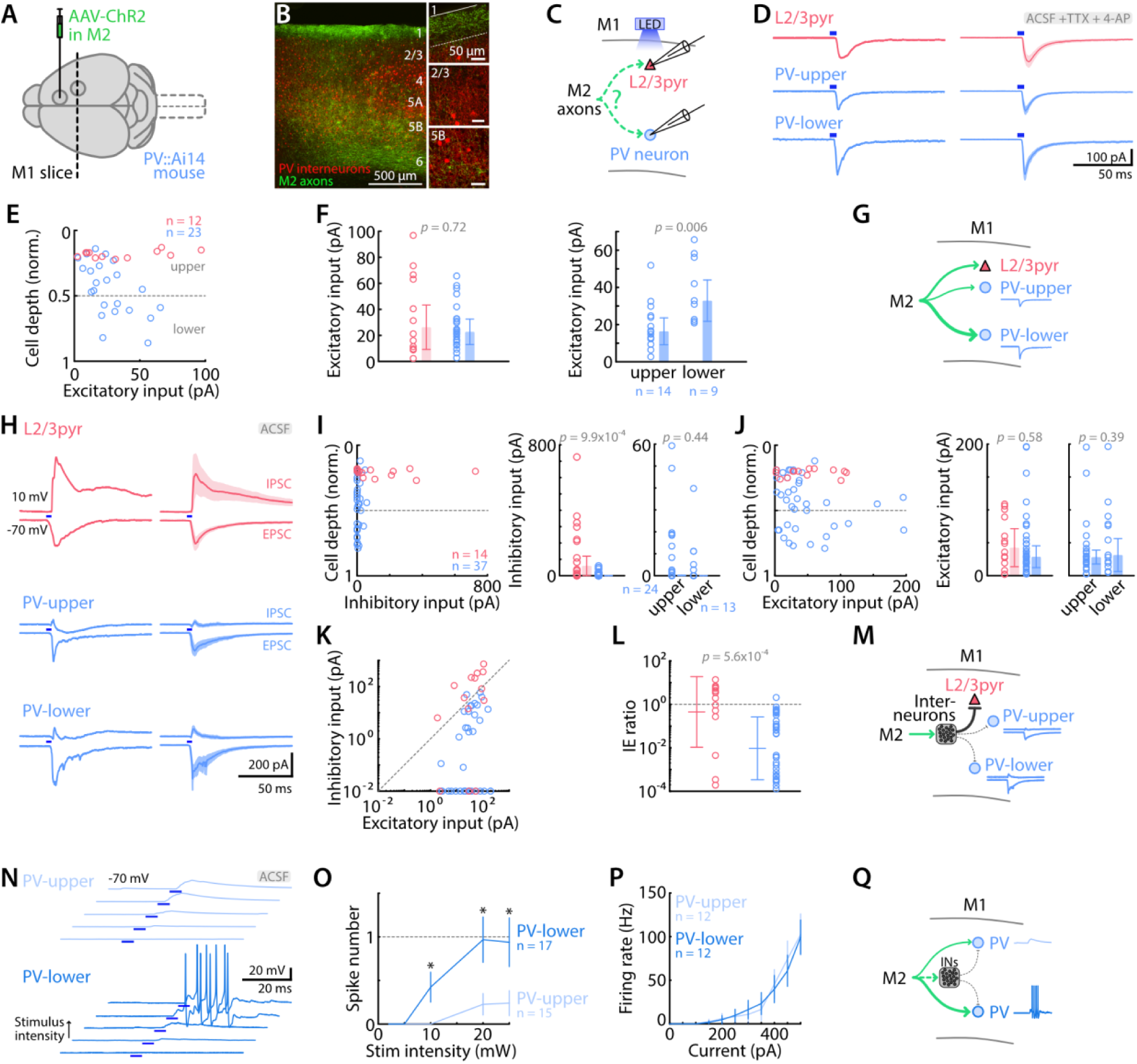
M2→M1 corticocortical input strongly excites PV interneurons without feed-forward inhibition. **A**, Anterograde AAV was injected in M2 to induce ChR2 expression in M1-projecting neurons in PV-Cre x Ai14 animals. **B,** Epifluorescence image showing M2 axons (green) and PV interneurons (red) along the cortical depth. Insets on the right show 60x magnification views of layer 1, 2/3, and 5B. **C,** M2 afferents were stimulated with blue light LED while M1 L2/3 pyramidal and PV interneurons responses were recorded. **D,** Example (left) and average (right, mean ± SEM) monosynaptic excitatory responses of L2/3 (red), and PV interneurons (blue) located in upper or lower layers, to photostimulation of M2 axons. Recordings were performed in artificial cerebrospinal fluid (ACSF) with TTX (1 µM), 4-AP (100 µM). Blue rectangles indicate LED stimulation. **E,** Excitatory input to L2/3 and PV interneurons in M1, calculated as the average response amplitude over 100 ms post-stimulus and plotted as a function of normalized cortical depth of the soma. Dashed line depicts the limit between upper and lower layers. **F,** Average excitatory input to L2/3 and PV interneurons in M1 (left) and amplitude difference between upper- and lower-layer PV interneurons responses (right). Bars show the median ± MAD. The two groups were compared using the Mann-Whitney U test. **G,** Schematic depiction of the monosynaptic excitatory connectivity pattern. **H,** Example (left) and average (right, mean ± SEM) excitatory and inhibitory responses of L2/3 neurons and PV interneurons located in upper or lower layers, to M2 stimulation (5 ms, 10 mW). Recordings were performed in plain artificial cerebrospinal fluid (ACSF). Blue rectangles indicate LED stimulation. **I,** Inhibitory input to L2/3 and PV interneurons in M1 plotted as a function of normalized cortical depth of the soma (left), average amplitude of L2/3 and PV interneurons (middle), and of upper and lower PV interneurons (right). Bars show the median ± MAD. **J,** Same for excitatory inputs. **K,** Amplitudes of excitatory and inhibitory responses recorded for each neuron. **L,** Ratio of inhibitory/excitatory (IE) input for each neuron (Mann-Whitney U test). **M,** Schematic depiction of the feedforward inhibitory connectivity pattern. **N,** Example traces of excitatory postsynaptic potentials evoked by M2 axons photostimulation with increasing light intensity (2, 5, 10, 20, 25 mW) in individual PV interneurons from upper and lower layers. **O,** Spike number as a function of stimulus intensity (mean ± SEM). Asterisks denote *p* value below 0.05 (one-way ANOVA). **P,** Frequency-current plot, showing the firing rate (number of action potentials per 1-s stimulus) as a function of current step amplitude. **Q,** Schematic depiction of the connectivity pattern.

Next, we recorded in control conditions (omitting TTX/4AP from the bath solution) to sample feedforward inhibitory responses. Stimulation of M2 axons evoked little or no inhibition in PV interneurons, contrasting with the relatively large inhibitory responses in L2/3 pyramidal neurons (0.08 ± 0.07 pA vs 59.7 ± 59.6 pA, *p* = 9.9×10^-4^; 37 PV interneurons, 14 L2/3 pyramidal neurons, from 8 animals). Inhibitory responses in PV interneurons were as low in upper layer as in lower layers (**Figure 4H, I**). Again, as found in the monosynaptic recordings above, the evoked EPSCs were of similar amplitude in L2/3 pyramidal neurons and PV interneurons (32.4 ± 22.2 pA vs 28.8 ± 16.6 pA, *p* = 0.58) (**Figure 4J**). While the average IE ratio of L2/3 pyramidal neurons was balanced, the ratio of PV interneurons favored excitation (0.46 (0.011 to 18.74) vs 0.009 (3.4×10^-4^ to 0.26), *p* = 5.6×10^-4^) (**Figure 4K, L**). These results thus demonstrate that the strong direct excitation of PV interneurons (reported above) is essentially unaccompanied by feedforward inhibition (**Figure 4M**).

As for corticospinal and L2/3 pyramidal neurons, we investigated how PV interneurons respond in current-clamp mode to M2 inputs. Similarly to corticospinal neurons, photostimulation of M2 axons with low intensity evoked EPSPs in deeper-layer PV interneurons and action potentials at higher intensities, sometimes in short trains. In contrast, responses in upper-layer PV interneurons showed lower spike probability with no or only single spikes (10 mW, *p* = 0.028, 20 mW, *p* = 0.020, 25 mW, *p* = 0.041, one-way ANOVA, 15 upper-layer and 17 lower-layer PV interneurons from 6 animals) (**Figure 4N, O**). The greater excitability of deeper-layer compared to upper-layer PV interneurons to M2 input was not ascribable to differences in intrinsic excitability, as the frequency-current relationships were similar (*p* = 0.92, one-way ANOVA, 12 upper layer and 12 lower-layer PV interneurons from 3 animals) (**Figure 4P**). These results thus indicate that PV interneurons, and those in deeper layers in particular, are readily recruited to fire action potentials by stimulation of M2 axons (**Figure 4Q**).

### M2 excitatory and inhibitory input to SST, VIP, and Ndnf neurons in M1

We performed the same experiment as above but with recordings targeted to other types of identified interneurons. We used mice expressing tdTomato in somatostatin (SST), vasoactive intestinal protein (VIP), or neuron-derived neurotrophic factor (Ndnf) interneurons (**Figure 5A**) (**Methods**). Interneurons in each line were again recorded across all layers (**Figure 5B**). For each recorded interneuron in each mouse line, we recorded both excitatory and inhibitory responses to M2 axon stimulation, and analyzed these as described above for the recorded excitatory neurons and PV interneurons.

**Figure 5.**
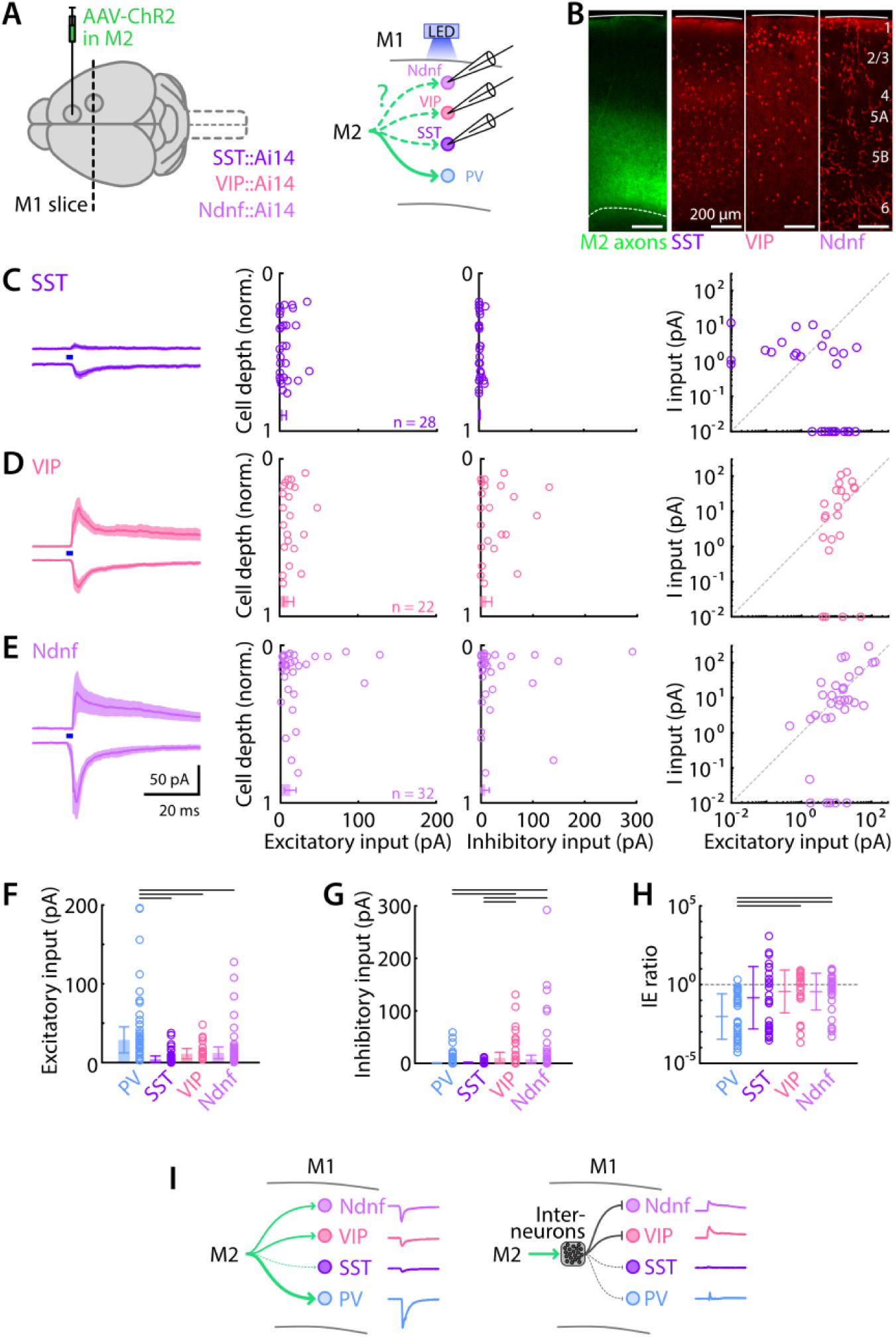
M2 excitatory and inhibitory input to SST, VIP, and Ndnf neurons in M1. **A**, Anterograde AAV-ChR2 was injected in M2 of mice with cell-type-specific labeling of interneurons, generated by crossing SST-, VIP-, and Ndnf-Cre driver lines with Ai14 tdTomato reporter mice (left). M2 axons were stimulated with blue light LED while M1 SST, VIP, or Ndnf interneurons responses were recorded (right). **B,** Epifluorescence image showing M2 axons (left - green) and cell distribution (right - red) for each interneuron type along the cortical depth. **C,** SST responses to M2 afferents stimulation, from left to right: group average traces (mean ± SEM) of evoked excitatory and inhibitory currents; strength of EPSCs and IPSCs as a function of normalized soma cortical depth; EPSC vs. IPSC input strength. **D,** Same, for VIP interneurons. **E,** Same, for Ndnf interneurons. **F,** Average values of excitation for each interneuron type. Black lines depict significant differences with *p* values below 0.5 (Kruskal-Wallis test with Dunn-Sidak correction for multiple comparisons). **G**, Same, for inhibition. **H,** Same, for IE ratios. **I,** Schematic depiction of the excitatory (left) and inhibitory (right) connectivity patterns.

Photostimulation of M2 axons evoked relatively weak excitatory and weak-to-absent inhibitory responses in SST interneurons (E: 4.25 ± 4.11 pA, I: 1.20 ± 1.19 pA, 28 neurons from 6 animals), with no obvious differences as a function of soma depth in the cortex (**Figure 5C**). For VIP interneurons (E: 11.3 ± 6.5 pA, I: 10.6 ± 10.6 pA, 22 neurons from 6 animals), both excitatory and inhibitory responses were common and of moderate amplitude on average, with no apparent depth dependence (**Figure 5D**). Similarly, Ndnf (E: 12.4 ± 7.4 pA, I: 7.77 ± 7.74 pA, 32 neurons from 6 animals) responses showed excitatory and inhibitory responses that were of moderate amplitude on average (**Figure 5E**).

Comparisons among all four interneuron classes showed that excitatory responses were greatest in PV interneurons (*p* values as follow: PV vs SST: 6×10^-8^, PV vs VIP: 0.015, PV vs Ndnf: 0.011, Kruskal-Wallis test with Dunn-Sidak correction for multiple comparisons) (**Figure 5F**). Inhibition was greater in VIP and Ndnf interneurons compared to PV and SST interneurons (PV vs VIP: 0.025, PV vs Ndnf: 0.011, SST vs VIP: 0.006, SST vs Ndnf: 0.002) (**Figure 5G**). Finally, PV interneurons’ inhibition-to-excitation ratio was more in favor of excitation than for the other interneuron types (PV vs SST: 0.008, PV vs VIP: 2×10^-4^, PV vs Ndnf 1×10^-4^) (**Figure 5H**).

These data thus demonstrate largely distinct patterns of excitation and inhibition evoked in these four classes of interneurons by photostimulation of M2 axons, with PV interneurons receiving the strongest excitation and weakest inhibition on average (**Figure 5I**).

### Local inhibitory inputs to M1 corticospinal neurons are strong from PV and SST, and weak from VIP and Ndnf interneurons

Having assessed M2 input to four classes of M1 interneurons, we next measured inhibitory input from the same interneuron classes to corticospinal neurons locally within M1. For this, we expressed ChR2 in the interneurons by crossing the driver Cre lines with the Ai32 reporter line, and used wide-field LED-based photostimulation to sample the evoked IPSCs in corticospinal neurons (**Figure 6A**). Of note, although some layer 5 excitatory neurons have been reported to express parvalbumin, we previously showed (Piña Novo et al., 2025) that for the mice used here (PV-Cre x Ai32), photostimulation-evoked responses are almost entirely GABAergic (i.e., not significantly contaminated by glutamatergic transmission).

**Figure 6.**
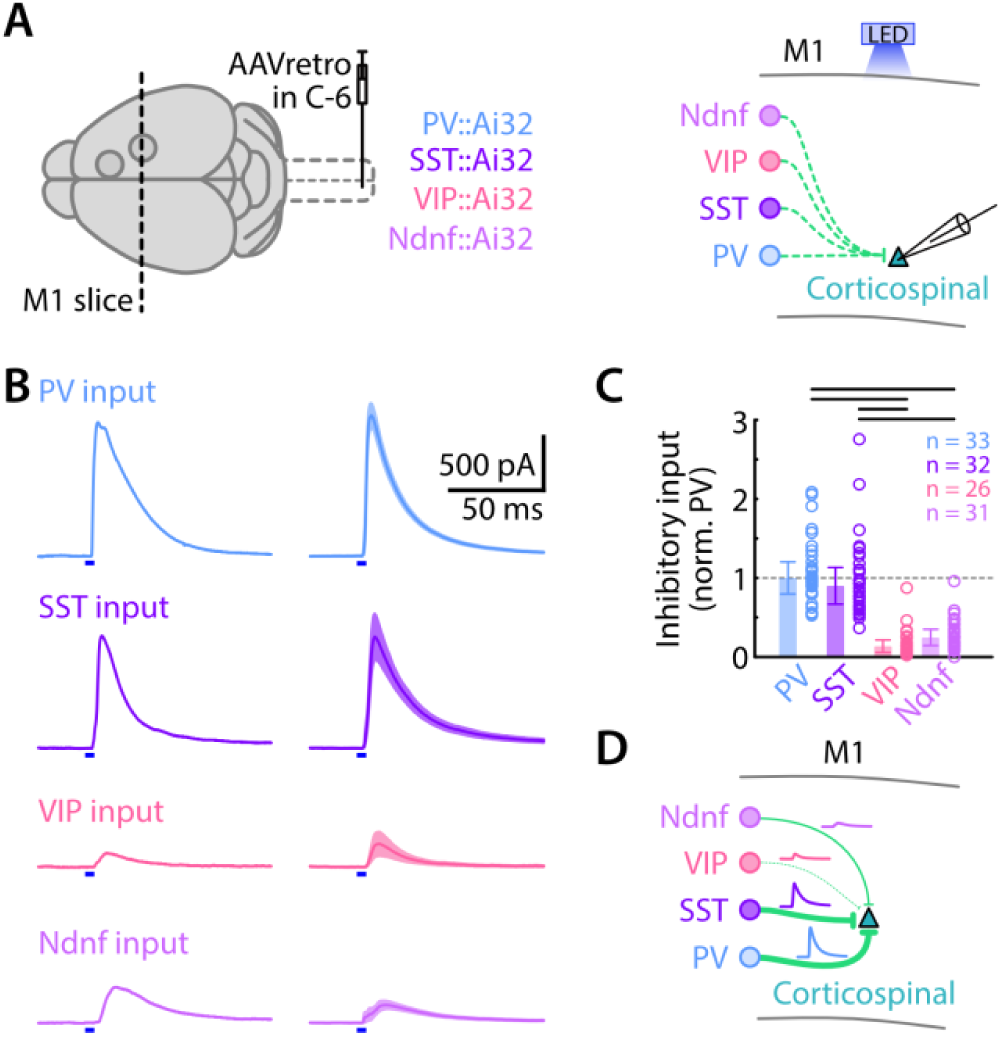
Local inhibitory inputs to M1 corticospinal neurons are strong from PV and SST and weak for VIP and Ndnf neurons. **A**, Corticospinal neurons were labeled by retrograde AAV injected in the spinal cord in mice crosses (Cre x Ai32) expressing ChR2 in the different interneuron lines: PV, SST, VIP, and Ndnf (left). Interneurons inputs were stimulated with blue light LED while M1 corticospinal neurons responses were recorded (right). **B,** Traces show evoked IPSCs in response to photostimulation of interneurons in M1 slices while recording from corticospinal. Example traces for individual neurons are shown along with the group average for each interneuron type stimulated (mean ± SEM) . **C,** Inhibitory input to corticospinal neurons, from each of the four interneuron types. Data are plotted as normalized input, relative to the group-average strength of PV input. Top lines indicate a *p* value below 0.05 (Kruskal-Wallis with Dunn-Sidak correction for multiple comparisons). **D,** Schematic depiction of the inhibitory connectivity from four classes of interneurons to L2/3 and corticospinal neurons. Lines thickness indicates inhibition strength.

Photostimulation of each interneuron type in each M1 evoked inhibitory responses in corticospinal neurons of varying amplitudes (**Figure 6B**). We plotted the relative strength of interneuron-specific inhibitory input for each. To pool data and compare across cell types, we normalized the inhibitory inputs by dividing by the average input from PV interneurons. The order of inhibitory input strength (from strongest to weakest) was: PV ≈ SST >> Ndnf > VIP (PV vs VIP: 1.8×10^-12^, PV vs Ndnf: 1.6×10^-8^, SST vs VIP: 2.1×10^-10^, SST vs Ndnf: 8.6×10^-7^, Kruskal-Wallis test + Dunn-Sidak corrected multiple comparisons; 33 PV from 4 animals, SST 32 SST from 6 animals, 26 VIP from 5 animals, 31 Ndnf from 4 animals) (**Figure 6C**). Thus, among these four interneuron classes, PV and SST interneurons were the strongest sources of local inhibitory inputs (**Figure 6D**).

### sCRACM-mapping of PV and SST inhibitory inputs to corticospinal neurons

With PV and SST interneurons identified as the cellular sources of local inhibition to corticospinal neurons, we also wanted to assess the subcellular organization of these connections, particularly as PV and SST inputs are expected to differentially target somata and dendrites, respectively. To do so, we prepared brain slices from mice with ChR2-expressing interneurons and tdTomato-expressing corticospinal neurons (**Figure 7A**).

**Figure 7.**
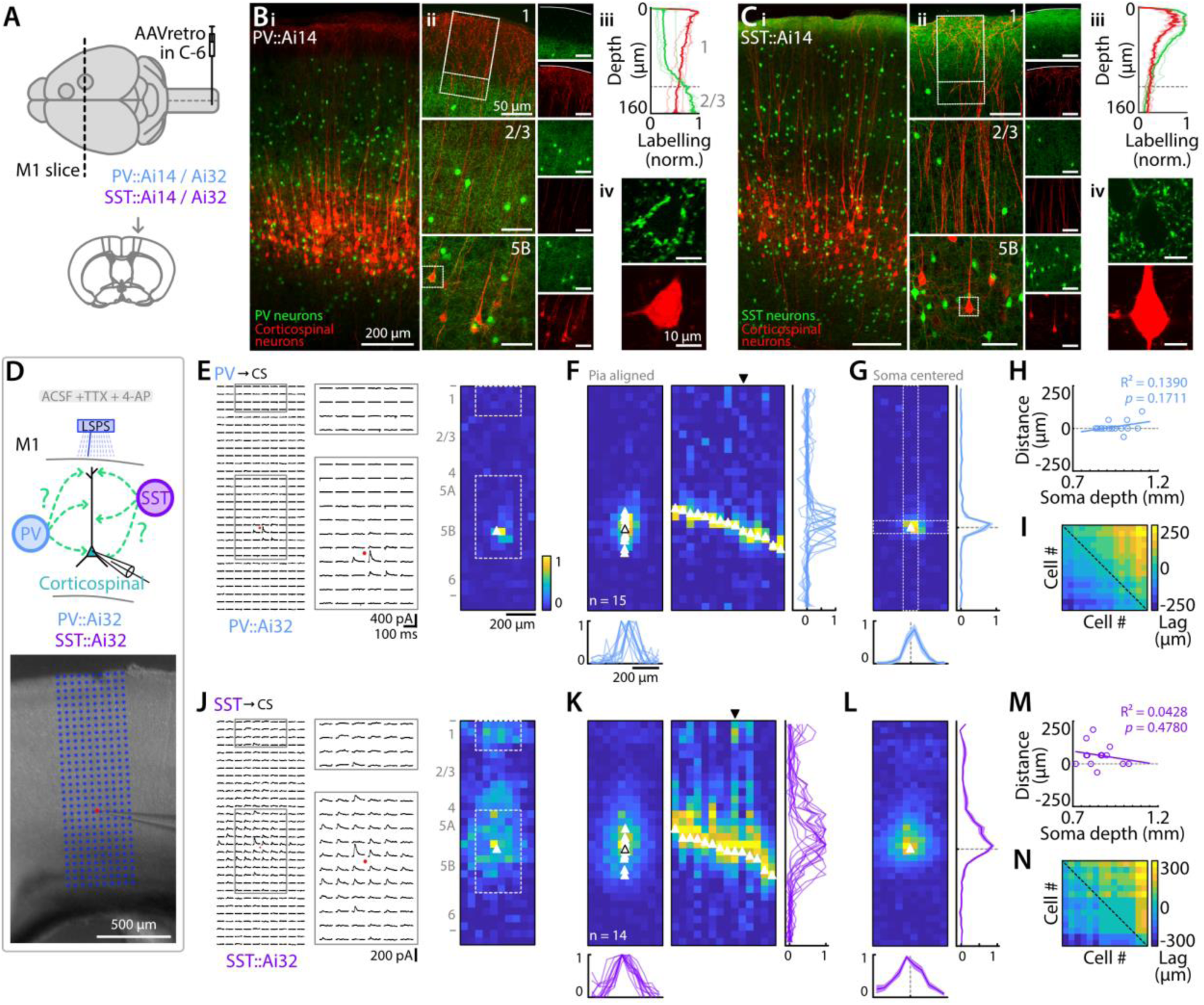
sCRACM-mapping of PV and SST inhibitory inputs to corticospinal neurons. **A**, Retrograde AAV-tdTomato was injected in the spinal cord in PV-Cre or SST-Cre mice crossed with either Ai14 mice (for imaging studies) or Ai32 mice (for optogenetic-electrophysiological studies). **B,** PV interneurons (green) and corticospinal neurons (red) are labeled in M1. Images show PV and corticospinal neurons labeling across all M1 cortical depth (i), higher magnification views of layer 1, 2/3, and 5B (ii, from top to bottom), plot profile of the labeling across the layer 1 / layer 2/3 border (designated area in ii) (iii), and a close-up view of a corticospinal neuron surrounded by PV terminals (designated by dashed square in ii) (iv). **C,** Same, for SST interneurons. **D,** Corticospinal neuron responses were sampled under sCRACM recording conditions (**Methods**) while laser-stimulating PV or SST inputs (top), and Image showing a M1 coronal slice with a patch pipette and an overlaying 10×30 stimulation grid horizontally centered on the pipette tip (i.e., neuron soma - red point) (bottom). **E,** Mapping of PV inputs on corticospinal neurons. Responses of an example neuron following laser stimulation in a 10×30 grid pattern (left). Soma is indicated by a red point. Enlarged views of perisomatic and apical regions are shown (middle). Heatmap of peak-normalized input strength measured from traces in the first panel (right). Soma is indicated by white triangle. **F,** Normalized responses averaged over neurons aligned on pia (left). Individual somata are indicated by white triangle and the average soma position is represented by an empty black triangle. Horizontal average projection of each neuron’s responses sorted by soma depth (middle) and corresponding horizontal and vertical profiles for each neuron (bottom and right). **G,** Normalized average response centered on soma with horizontal (left) and horizontal and vertical average projections (2 pixels width centered on soma) of input strength centered on soma (dashed line, mean ± SEM) (bottom and right). **H,** Distance from the peak value of each neuron’s vertical profile to its soma, as a function of soma depth in the cortex. Line: linear regression. **I,** Cross correlation analysis. Each element in the pseudocolored matrix represents the lag (distance) at the peak value of the cross correlation for the vertical profiles of one pair of neurons’ input maps (see **Methods**). Dashed line indicates main diagonal. **J-N,** Same, for SST inputs.

Anatomically, PV somata were distributed in all layers of M1 except for layer 1. Labeling of dendrites and axons was observed in all layers except for layer 1 (**Figure 7Bi-iii**). Corticospinal neurons cell bodies and proximal dendrites were surrounded by PV somata and dendritic and axonal processes (**Figure 7Bi, ii**), with dense labeling surrounding corticospinal somata (**Figure 7Biv**). The SST interneurons were also distributed in all layers of M1 except for layer 1, but projected a higher density of axonal terminations in layer 1, such that corticospinal neurons apical dendrites were overlapping with SST axons (**Figure 7Ci-iii**). Unlike PV labeling, SST labeling was not clustered around corticospinal somata (**Figure 7Civ**).

We mapped the distribution of inhibitory inputs to corticospinal dendrites using an optogenetic LSPS-based method, sCRACM, with conditions set to isolate monosynaptic inputs (TTX/4AP) (**Methods**). The stimulus array for mapping was oriented to cover the full cortical depth, aligned vertically to the pia surface and horizontally to the corticospinal neuron’s soma (**Figure 7D**). Maps of PV input to corticospinal dendrites showed that IPSPs were generally evoked only in the immediate vicinity of the soma (**Figure 7E**). This pattern is consistent with expectations given that PV axons are soma-targeting (Hu et al., 2014), and with previous LSPS maps of perisomatic inputs from PV to layer 5 pyramidal neurons in mouse cortex (Brill et al., 2016). For corticospinal neurons located at different cortical depths from the pia (and thus within L5B), the small zone of inhibitory input was consistently over the soma, independent of cortical depth; i.e., the input pattern was soma-centric, not cortex-centric (15 neurons from 4 animals) (**Figure 7F, H**). Consistent with this, averaging the maps of multiple neurons showed a vertically smeared zone of input when maps were pia-aligned, but a sharply peri-somatic zone of input when maps were soma-aligned. Similarly, when we converted each map to a vertical profile and then calculated the cross-correlation of the group of profiles (**Methods**), this also showed that map structure correlated more with postsynaptic somatodendritic domain structure than with presynaptic axonal laminar structure (**Figure 7I**). In other words, maps of different neurons were more correlated with each other than they were with cortical laminar morphology, again most consistent with a postsynaptic soma-centric organization of inputs.

Maps of SST input to corticospinal dendrites (14 neurons from 5 animals) (**Figure 7J-N**) showed that IPSCs were evoked across a broader somatodendritic territory surrounding the soma. This pattern is largely consistent with expectations given that SST axons are dendrite-targeting (Urban-Ciecko and Barth, 2016). Again, the pattern was mainly soma-centric, as seen in the soma-centered average map and the correlation analysis. Inputs to the distal dendrites in the apical tufts were observed but were generally relatively weak, attributable at least in part to the long electrotonic distance.

These results confirm, using a functional method for mapping synaptic inputs across dendritic arbors, that PV and SST interneurons target corticospinal neurons in a soma-centered manner, but differentially targeting the soma and dendritic compartments, respectively.

### LSPS mapping of local sources of PV and SST inhibitory input to corticospinal neurons

The sCRACM results show which domains of corticospinal dendritic arbors receive inputs, but not where in the local circuit those inputs arise from. To address this question, we used the same Cre lines as previously but crossed them with Ai203 mice, which express soma-targeted channelrhodopsin ChroME (with GCaMP7s and mRuby) in cells containing Cre (Bounds et al., 2023). This enabled soma-targeted photostimulation and thus LSPS mapping of local sources of PV and SST input.

First, we confirmed the soma-targeted photoexcitability of st-ChroME-expressing interneurons by recording excitation profiles. For this, we recorded from mRuby-expressing PV or SST interneurons in the presence of NBQX (10 µM) and gabazine (10 µM), to allow action potentials but block excitatory and inhibitory synaptic transmission. As expected (Bounds et al., 2023), in the subset of these neurons that expressed ChroME, excitation profiles showed that sites of photoexcitability were restricted to the perisomatic region, for both PV (3 neurons from 2 animals) and SST (7 neurons from 2 animals) interneurons (**Figure 8A, B**).

**Figure 8.**
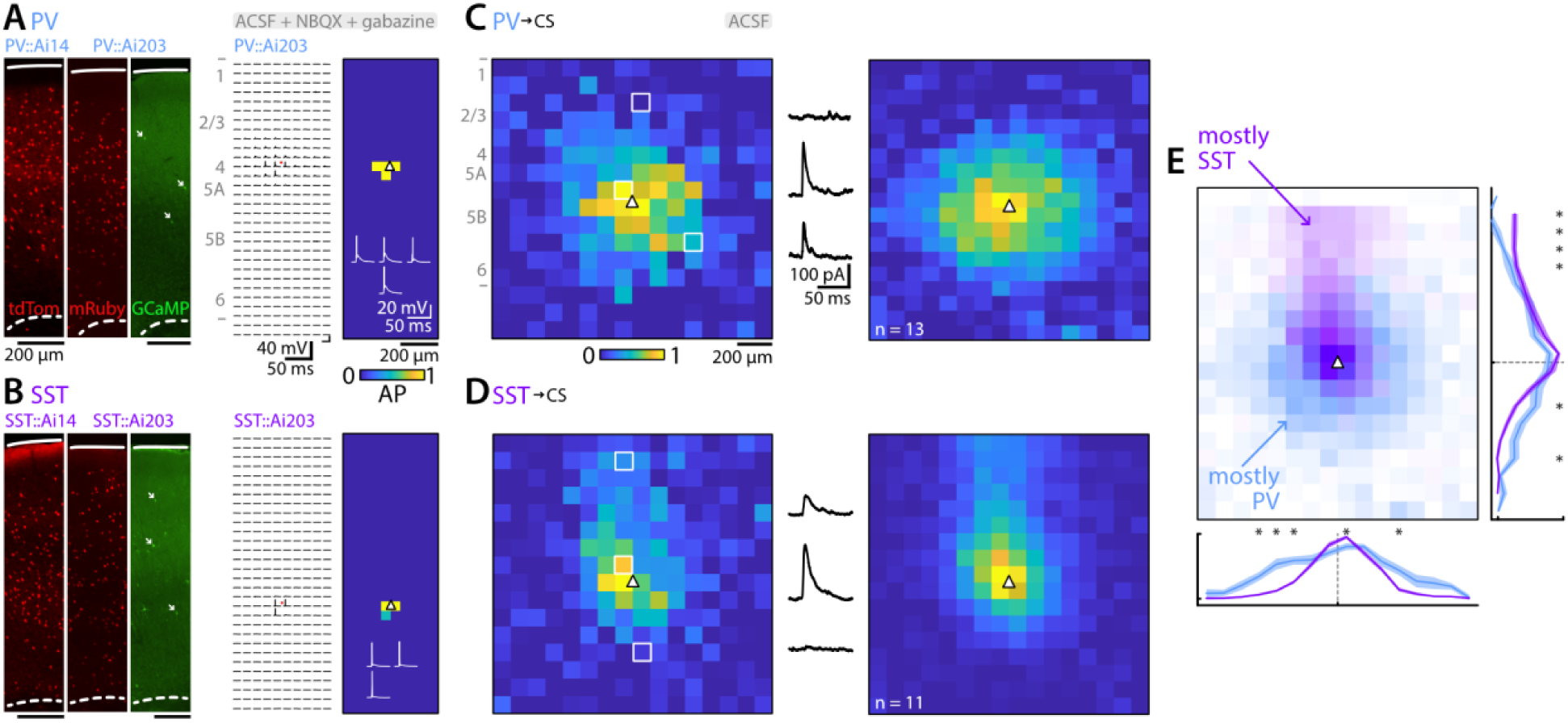
LSPS mapping of local sources of PV and SST inhibitory input to corticospinal neurons. **A**, Excitation profile of a PV interneuron expressing ChR2 in PV x Ai203 mice. Left: example images showing labeling of interneurons in PV-Cre mice crossed with either Ai14 mice (left image) or Ai203 mice (middle and right images). In the Ai203 cross, Cre drives expression of nuclear-targeted mRuby in all neurons (middle image) and of the opsin-GCaMP fusion construct in a subset of these (right image; arrows mark several examples of labeled somata). Middle: example of action potentials evoked for somatic stimulation. Right: heatmap of AP number. PV soma indicated by white triangle. Traces with evoked action potentials are also shown. **B,** Same for an SST interneuron. **C,** Maps of local sources of PV inhibitory input to corticospinal neurons. Left: example of single neuron (normalized input) with evoked IPSC traces at different stimuli locations (indicated by white squares). Right: soma-centered average. Average corticospinal soma position indicated by white triangle. **D,** Same for SST interneurons. **E,** Comparison of PV and SST input maps to corticospinal neurons and vertical and horizontal profiles. Asterisks indicate a p value below 0.05 (Mann-Whitney U test).

Then, using plain ACSF (omitting TTX, 4AP, and synaptic blockers), we recorded from labeled postsynaptic corticospinal neurons while mapping inhibitory inputs evoked from PV (13 neurons from 6 animals) or SST (11 neurons from 3 animals) interneurons using LSPS. Although only a subset of mRuby-labeled neurons in these crosses are expected to also express the opsin (Bounds et al., 2023), input maps for both the PV and SST lines showed robust inputs, which arose from local regions surrounding the corticospinal neurons (**Figure 8C, D**). However, the zone of PV input was more extended horizontally (i.e., wider), whereas the zone of SST input was more extended vertically toward the pia (taller) (**Figure 8E**).

These results demonstrate that, in addition to targeting different postsynaptic domains, inhibitory inputs from PV and SST interneurons to corticospinal neurons arise from presynaptic interneurons located in partly overlapping, partly distinct local regions.

## DISCUSSION

Using slice-based optogenetic and electrophysiological methods to dissect cell-type-specific circuits, we assessed synaptic connectivity linking presynaptic M2 axons to major classes of postsynaptic M1 excitatory and inhibitory neurons. The results identify basic elements of these forelimb-related premotor-to-motor corticocortical circuits in the mouse (**Figure 9**).

**Figure 9.**
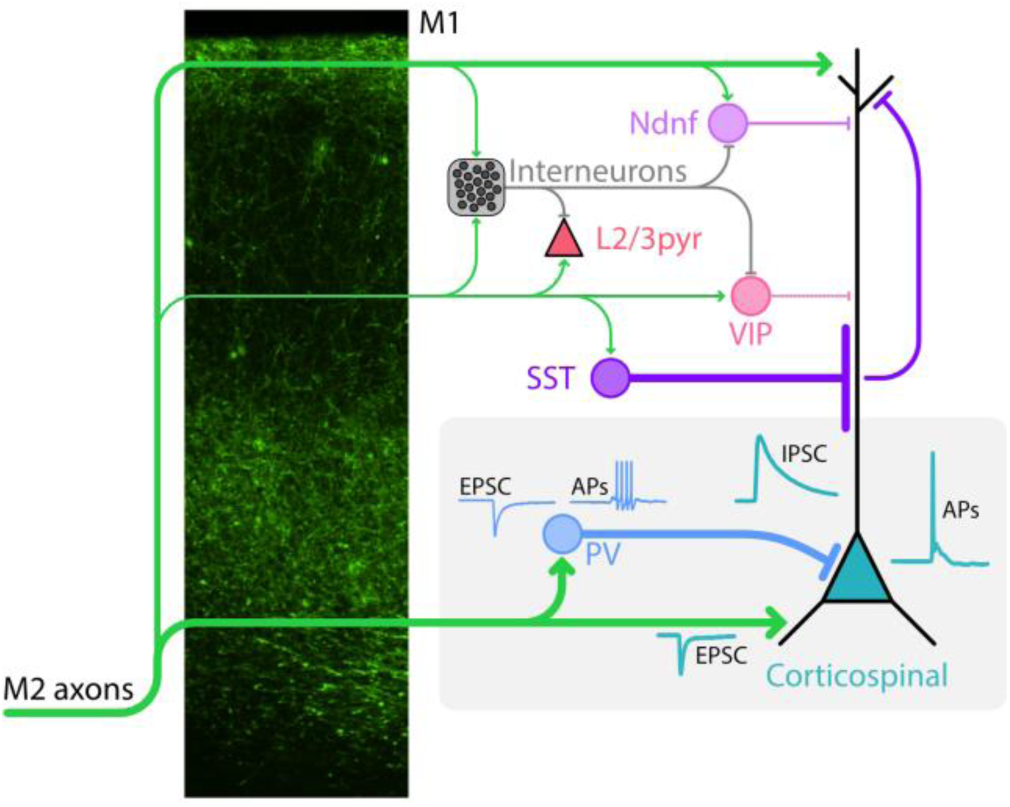
Schematic summary. The schematic depicts the main results of responses recorded in different M1 neurons during photostimulation of the axons of either M2→M1 corticocortical projections or local interneurons. The fluorescence image shows the laminar profile of labeled M2 axons in M1. The grey boxed region highlights the strongest pathways identified, including direct excitation and PV-mediated feedforward inhibition of M1 corticospinal neurons.

Direct excitation of corticospinal neurons is a prominent feature of these M2→M1 circuits. Anatomically, corticocortical axons that arise primarily from layer 5A neurons of M2 ramify in M1 mainly in a broad band across deeper layers, plus a narrow band in layer 1. The laminar profile of these afferent axons is essentially bookmatched to the laminar profile of M1 corticospinal dendrites, a pattern of axo-dendritic overlap that suggests functional specialization of M2→M1 circuits for direct excitation of corticospinal neurons. Indeed, subcellular mapping previously detected M2 excitatory inputs to the apical and perisomatic/basal dendritic subarbors of corticospinal neurons in forelimb M1 (Suter and Shepherd, 2015), similar to observations in M1-to-S1 circuits (Petreanu et al., 2009). Here, we confirmed the strong monosynaptic excitatory connectivity from M2 axons to M1 corticospinal neurons and show both the ability of these connections to readily drive action potentials in corticospinal neurons and their relative strength compared to inputs to L2/3 pyramidal neurons. We also show that these L2/3 pyramidal neurons sample inputs both in layer 1 and layer 2/3. From a broader perspective, because top-down projections generally ramify in a similar L1+L5/6 pattern (D’Souza and Burkhalter, 2017; Harris et al., 2019; Liu et al., 2024) and pyramidal tract (PT) neurons are ubiquitous across cortex (Harris and Shepherd, 2015), we speculate that the connectivity observed here exemplifies a generic property of top-down pathways; i.e., higher-order upstream areas have direct synaptic influence on PT neuron excitability and thus on cortical output in lower-order downstream areas. In contrast, in bottom-up pathways, afferent projections do not appear specialized to match so precisely the dendritic arbors of a single class of excitatory neurons (Mao et al., 2011; Harris et al., 2019).

The M2 excitation of M1 corticospinal neurons was accompanied by strong feedforward inhibition, ascribable primarily to PV interneurons as they receive direct and potent excitation from M2 (which is unaccompanied by feedforward inhibition, and readily drives action potentials), and they in turn deliver strong soma-targeting GABAergic input to corticospinal neurons. These results align with and complement prior findings of strong PV-mediated inhibition of PT-type neurons, which can be driven by long-range corticocortical inputs and may be critical for dampening excitation (Anastasiades et al., 2018) as well as gamma rhythmogenesis (Kawaguchi et al., 2019). Among other inhibitory neurons in M1, both excitatory and inhibitory inputs from M2 axons were relatively weak to SST interneurons, and stronger to VIP and Ndnf neurons. Locally within M1, monosynaptic inhibition to corticospinal neurons was strong from SST as well as PV interneurons, while inputs from VIP and Ndnf were relatively weak. These findings largely agree with prior results from other areas and pathways. Using Cre-driver mouse lines to label GABAergic cell classes and targeting recordings to labeled interneurons distributed across the cortical depth, we detected one intra-class difference, finding greater excitation of PV interneurons in lower compared to upper layers. As each of the top-level classes of interneurons studied here contains multiple subtypes (Schuman et al., 2019; Scala et al., 2021; Wu et al., 2023), further work using more selective methods (Tasic and Fishell, 2025) may reveal subtype-specific circuit properties.

The topography of sCRACM-mapped inhibitory inputs to corticospinal neurons generally matched expectations based on the soma- and dendrite-targeting properties of PV and SST interneuron inputs to pyramidal neurons. In the SST maps, although inputs could be detected throughout the dendritic arbor including the apical tuft in layer 1, responses were much stronger around the soma and weaker distally. This likely reflects biophysical and morphological factors: (i) strong electronic attenuation along dendrites, causing a distance-dependent drop in response amplitudes (Williams and Mitchell, 2008), and (ii) highest concentration (length density) of dendrites around the soma (Suter and Shepherd, 2015). Similar distance-dependent effects have been observed in maps of glutamatergic responses across corticospinal neuron dendritic arbors (Anderson et al., 2010; Sheets et al., 2011; Hooks et al., 2013; Suter and Shepherd, 2015), and considered in the context of sCRACM map interpretation (Petreanu et al., 2009).

On the presynaptic side, local PV and SST inputs to corticospinal neurons arose from partly overlapping perisomatic domains, wider for PV inputs and taller for SST inputs. This suggests that mainly deeper- and not upper-layer PV neurons provide disynaptic inhibition from M2 axons to corticospinal neurons; in contrast, the inhibitory input via SST neurons, although much smaller compared to PV neurons, may arise from SST neurons across multiple layers. Interestingly, these patterns partly resemble those of local excitatory input to PV and SST interneurons in layer 5A/B of mouse motor cortex, with inputs to PV arising mostly from deeper layers, and inputs to SST arising mainly from upper layers (Apicella et al., 2012).

Our characterizations of forelimb-related M2→M1 circuits in the mouse build on prior results for these and related motor/frontal corticocortical circuits (Rouiller et al., 1993; Hira et al., 2013; Hooks et al., 2013; Ueta et al., 2014; Harris et al., 2019). The vibrissal M1→S1 has served as a model top-down pathway, with its excitatory axons targeting multiple cell types including infragranular pyramidal, VIP, and PV interneurons (Petreanu et al., 2009; Lee et al., 2013; Kinnischtzke et al., 2014, 2016; Naskar et al., 2021; Kim et al., 2024). In S2→S1 but not M1→S1 circuits the afferent axons preferentially target PV over VIP interneurons (Naskar et al., 2021), and hindlimb M2→S1 circuits show similar top-down circuit organization, biased toward infragranular pyramidal and PV interneurons (Manita et al., 2015; Atsumi et al., 2023). Our finding of stronger M2→M1 excitation of PV than VIP interneurons appears to contrast with a recent study of corticocortical pathways in the visual system (Ma et al., 2021), which concluded that top-down pathways generally target PV interneurons while bottom-up pathways target VIP interneurons; however, that study focused on neurons in layer 2/3, which here received less excitation than deeper-layer PV interneurons.

The circuit organization of forelimb M2→M1 differs from that of other corticocortical pathways to M1. In particular, the hand/forelimb S1→M1 pathway, the main bottom-up pathway conveying somatosensory signals along the transcortical loop, mainly targets M1 pyramidal neurons in layer 2/3, with little or no direct excitatory input to corticospinal neurons (Yamawaki et al., 2021; Piña Novo et al., 2025). In counterpart vibrissal pathways, whisker S1→M1 inputs target upper-layer pyramidal and PV interneurons, and middle-layer SST interneurons (Mao et al., 2011; Okoro et al., 2022; Goz and Hooks, 2023). In S2→M1, hierarchically a more “lateral” type of pathway, excitatory input favors upper layers but with moderate input to corticospinal neurons (Suter and Shepherd, 2015). Individually, the multiple input pathways to M1 typically each innervate a particular subset of cell types and layers; collectively, they appear to innervate most if not all major cell types across layers (Hooks et al., 2013; Harris and Shepherd, 2015).

This study used slice-based methods and focused primarily on static rather than dynamic synaptic connectivity, using single rather than repetitive photostimuli. An area for further investigation is the short-term plasticity at each of the many cell-type-specific connections identified here (Kamalova et al., 2024). Short-term dynamics of corticocortical synapses to corticospinal neurons may facilitate (Kim et al., 2024), but those of feedforward PV and SST circuits may boost net inhibition (Galarreta and Hestrin, 1998; Silberberg and Markram, 2007). Further studies should characterize in vivo spiking dynamics. In the hand/forelimb-related S1→M1 pathway, close correspondence was found between slice-based circuit connectivity and in vivo spiking dynamics (Yamawaki et al., 2021; Piña Novo et al., 2025). The combination of linear array electrophysiology and optogenetic manipulations can facilitate mechanistic investigation of the cell-type-specific connections characterized here. Such approaches can identify the circuit-level mechanisms by which dynamic activity along top-down pathways interacts with bottom-up signaling, to route or otherwise modulate motor cortical output (Zagha, 2020). More generally, detailed knowledge about M2-M1-S1 circuits makes this a model system for elucidating the multi-region interactions that mediate integration of cognitive, motor, and sensory information (Shepherd and Yamawaki, 2021) through application of cell-type-specific tools for circuit analysis (Adesnik and Naka, 2018; Luo et al., 2018; Tasic and Fishell, 2025).

## Acknowledgments

This work was supported by National Institutes of Health–National Institute of Neurological Disorders and Stroke Grant R37NS061963 to G.M.G.S. Imaging work was performed at Northwestern University’s Center for Advanced Microscopy (RRID: SCR_020996) generously supported by NCI CCSG P30 CA060553, awarded to the Robert H. Lurie Comprehensive Cancer Center. We thank Megan Martin for technical assistance and John M. Barrett, Mang Gao, and Daniela Piña Novo for comments and advice.

## Author contributions

L.R. and G.M.G.S. designed research; L.R., R.F., and M.B. performed research; L.R., R.F., and M.B. analyzed data; L.R. and G.M.G.S. wrote the paper.

